# Salivary metabolomics in the family environment: A large-scale study investigating oral metabolomes in children and their parental caregivers

**DOI:** 10.1101/2024.02.21.581494

**Authors:** Jason A. Rothman, Hillary L. Piccerillo, Jenna L. Riis, Douglas A. Granger, Elizabeth A. Thomas, Katrine L. Whiteson

**Affiliations:** Department of Molecular Biology and Biochemistry, University of California, Irvine, Irvine, CA, 92697; Institute for Interdisciplinary Salivary Bioscience Research, University of California, Irvine, Irvine, CA, USA; Department of Health and Kinesiology, University of Illinois at Urbana-Champaign, Urbana, IL, USA; Department of Psychological Science, University of California, Irvine, Irvine, CA, USA; Department of Pediatrics Johns Hopkins University School of Medicine, Baltimore, MD, USA; Department of Neurobiology and Behavior, University of California, Irvine, Irvine, CA, USA

## Abstract

Human metabolism is complex and dynamic, and is impacted by genetics, diet, health, and countless inputs from the environment. Beyond the genetics shared by family members, cohabitation leads to shared microbial and environmental exposures. Furthermore, metabolism is affected by factors such as inflammation, environmental tobacco smoke (ETS) exposure, metabolic regulation, and exposure to heavy metals.

Metabolomics represents a useful analytical method to assay the metabolism of individuals to find potential biomarkers for metabolic conditions that may not be phenotypically obvious or represent unknown physiological processes. As such, we applied untargeted LC-MS metabolomics to archived saliva samples from a racially diverse group of elementary school-aged children and their caregivers collected during the “90-month” assessment of the Family Life Project. We assayed a total of 1,425 saliva samples of which 1,344 were paired into 672 caregiver/child dyads. We compared the metabolomes of children (N = 719) and caregivers (N = 706) within and between homes, performed population-wide “metabotype” analyses, and measured associations between metabolites and salivary biomeasures of inflammation, antioxidant potential, ETS exposure, metabolic regulation, and heavy metals.

Dyadic analyses revealed that children and their caregivers have largely similar salivary metabolomes. Although there were differences between the dyads at the individual levels of analysis, dyad explained most (62%) of the metabolome variation. At a population level of analysis, our data clustered into two large groups, indicating that people likely share most of their metabolomes, but that there are distinct “metabotypes” across large sample sets. Lastly, individual differences in several metabolites – which were putative oxidative damage-associated or pathological markers – were significantly correlated with salivary measures indexing inflammation, antioxidant potential, ETS exposure, metabolic regulation, and heavy metals. Implications of the effects of family environment on metabolomic variation at the population, dyadic, and individual levels of analyses for health and human development are discussed.

## Introduction

Metabolomics allows for the assay and quantification of thousands of chemical compounds in a biological sample (1–5). In humans, metabolomics correlates with the overall metabolism of biochemical pathways in not only the individual, but also microbes that inhabit specific areas of the body (1,6). Previous studies have shown that metabolomics is a useful tool that has the potential to find biomarkers for diseases and stress, generally through analyzing bodily fluids from subjects (7–9). One such sample specimen is saliva - a complex matrix of chemical compounds, proteins, and cells important in lubricating the oral cavity, digesting food, and protecting against infection (3,10,11). As a minimally-invasive biofluid to collect, saliva offers a straightforward way to assay metabolites, proteins, and xenobiotic compounds to find biomarkers for disease or environmental stress exposure (12–14). Aside from host metabolites, saliva also notably contains microbially-produced compounds and provides a medium to associate host and microbial relationships (1,6,15). In this study, we concentrate on inter/intrafamily relationships of the salivary metabolome, and associations between this metabolome and biomeasures relating to environmental tobacco smoke (ETS) exposure (the nicotine breakdown product cotinine) (16), antioxidant potential (uric acid) (12), inflammation (C-Reactive Protein; CRP) (17), and metabolic regulation (adiponectin) (13), along with a selection of metals (chromium, copper, lithium, manganese, and zinc) (14).

Family members are known to share similar oral health attributes, living environments, and overall diets. Likely due to this close social ecology of the home, several studies have shown that the oral microbiome is more similar within cohabitating family members than those outside of the home (18–21). Perhaps more surprisingly, the overall metabolomes of parental caregivers and their children are more similar than nonfamily individuals, and several metabolites are strongly correlated between family members (22–26). A well-studied example of these associations is within breastfeeding mothers: Metabolites that mothers produce affect the metabolomes and health of their infants through breastfeeding, and mothers’ environmental chemical exposures potentially affect the growing fetuses in utero through placental exposure (27,28). While the concept of interfamilial metabolic similarity is not novel, many studies concentrate on well-established health biomarkers such as cholesterol, amino acids, and hormones. However, due to the targeted nature of these analytical methods, likely do not capture overall metabolic relationships within families, and may miss currently-unknown biomarkers of oral health (8,29). Much like the oral microbiome (1,18), the salivary metabolome can be categorized into multiple “ecotypes” of metabolites that may correspond to oral dysbiosis, altered biochemical profiles, or baseline metabolism (1,9,30), but exactly how widespread this clustering is across large populations remains largely unknown.

The human oral cavity contains a multitude of chemicals derived from host biofluids, and as mentioned above, allows for investigations into the environment in which people live through salivary analyses (10,14). For example, kit-based and spectrometry-based methods can detect the presence of analytes involved in nicotine metabolism (i.e. cotinine), drugs and medications, xenobiotics, and metals (14,31,32). Similarly, through untargeted metabolomics, we may be able to discover novel biomarkers of oral dysbiosis (or conversely, health) that provide new diagnostic tools and establish baselines measurements of chemicals within saliva (1,3,8,15). Families living in rural or lower-socioeconomic areas are often exposed to high pollution burdens (14,33,34), which likely contributes to chronic stress. Furthermore, individuals exposed to heavy metals are known to have higher metabolites associated with oxidative stress and damage (reviewed in Bonvallot 2018 (35). As part of further understanding the effects that environmental stressors have on children (36), we chose to analyze salivary metals content and associate them with our untargeted metabolomes. These elements serve as reliable indicators of exposure, and we seek to uncover potential biomarkers for metals exposure and add to the growing field of “exposomics” (37–39).

Given the importance of understanding human metabolism and how it relates to families, we investigated the salivary metabolome in the context of children and their caregivers as part of the Family Life Project (FLP), a large-scale prospective longitudinal study (36). For example, if caregivers are smoking - with negative impacts on oral and other health measures - are these markers shared with children who are not smoking, but share the same household? We asked several questions of our data: First, does the salivary metabolome differ between children and adults (caregivers, henceforth), and is there metabolic concordance within families? Second, does the salivary metabolome associate with biomeasures of ETS exposure, antioxidant potential, metabolic regulation, or inflammation, and are there chemicals in saliva that may serve as additional biomarkers for the above-mentioned biomeasures? Lastly, are there associations between salivary metals burden and metabolites, and might these associations indicate potential negative health outcomes?

## Materials and methods

### Study participants

The Family Life Project (FLP) is a longitudinal study of families residing in Pennsylvania or North Carolina. FLP began in 2003-2004, when a representative sample of 1,292 children whose families resided in the target communities at the time the mothers gave birth were recruited and enrolled. Detailed descriptions of the sampling and recruitment procedures are available in Vernon-Feagans, Cox, and the FLP Key Investigators, 2013 (36). Briefly, families with a child born between September 2003 and August 2004 were recruited from hospitals, and multiple subsequent FLP study visits have been conducted with consented participants from children aged 2-months to age 20 years. The current analyses focused on a subset of data collected at the child’s 90-month at-home follow-up visit, where children and their primary caregivers provided unstimulated, resting/baseline saliva samples via passive drool collection. These saliva samples were assayed and archived in -80°C freezers, and biospecimens with adequate saliva remaining for metabolome analysis were examined in this study. Out of the total FLP sample, we assessed 1,425 saliva samples (child N for this subsample = 719; female = 353, male = 366; age = 79-100 months [average = 87 months], caregiver N = 706; female = 692, male = 27; age = 22-65 years [average = 34 years) of which 1,344 were paired into 672 caregiver/child dyads. Procedures for this study were run under the Institutional Review Boards of the University of North Carolina (IRB # 07-0646 and 16-2751) and New York University (IRB # IRB-FY2017-69) using deidentified data. Sample IDs were further randomized prior to analysis and reporting.

### Saliva sample handling for biomeasure analyses

All salivary biomarker analyses were conducted at the UC Irvine Institute for Interdisciplinary Salivary Bioscience Research (IISBR), where samples were stored at -80⁰C as previously described (18). Briefly, we thawed the samples, vortexed, then centrifuged samples for 15 minutes at 3500 RPM, then analyzed the supernatant for adiponectin, CRP, cotinine, and uric acid concentrations as follows below. We note that only children’s samples were assayed for all biomeasures, while caregiver samples were subject to cotinine analysis only. Samples numbers for each analyte in children were: adiponectin (N = 701), CRP (N = 616), cotinine (N = 714), and uric acid (N = 630), while caregivers’ cotinine was measured in (N = 672) samples. *Adiponectin, C-Reactive Protein, Cotinine, and Uric acid assays:*

We analyzed salivary adiponectin with the Human Adiponectin MSD assay kit (Meso Scale Discovery, Rockville, MD, USA). We diluted samples five-fold, then assayed following the manufacturer’s supplied protocol using a four-log standard curve and read the sample concentrations on a Meso Quickplex SQ120 spectrophotometer (Meso Scale Discovery, Rockville, MD). Concentrations were derived using MSD Discovery Workbench software v4.0 using curve fit models, with an assay range of sensitivity of 0.06 to 1000 ng/mL.

We assayed CRP in duplicate with the Human CRP V-Plex MSD Multi-spot Assay kit (Meso Scale Discovery, Rockville, MD, USA). We diluted samples five- or ten-fold, then assayed following the manufacturer’s protocol using a modified four-log standard and read the sample concentrations on a Meso Quickplex SQ120 spectrophotometer. The assay range sensitivity was 1.33 – 46,600 pg/mL CRP.

We measured salivary cotinine concentrations in both children and caregivers using the Salimetrics Salivary Cotinine ELIZA kit (Salimetrics, Carlsbad, CA, USA) following the manufacturer’s protocol. We analyzed 20 uL of sample in duplicate by incubating samples with kit reagents for 90 minutes with shaking at 37°C, and diluted samples 10-fold if subjects reported nicotine a Meso Quickplex SQ120 spectrophotometer use. As above, we washed the plates then added TMB followed by room temperature incubation for 30 minutes in the dark. We added 2M sulfuric acid and read the results on a Meso Quickplex SQ120 spectrophotometer and computed a standard curve using a four-parameter non-linear regression curve fit. The assay range of sensitivity was 0.15 to 200 ng/mL for neat saliva and 1.5 to 2000 ng/mL for 10-fold diluted saliva, and we substituted values of ½ the lower limit of measurement for each sample under the lowest reliable measurement for 316 samples (N = 138 caregivers and N = 178 children).

We analyzed uric acid with the Salimetrics Salivary Uric Acid Assay kit (Salimetrics, Carlsbad, CA, USA). We mixed 10 uL of sample with 190 uL of uric acid reagent in duplicate following the manufacturer’s protocol, then measured the results on a PowerWave HT spectrophotometer (BioTek/Agilent Technologies, Santa Clara, CA). The uric acid assay had a range of sensitivity from 0.07 – 20 mg/dL.

### Salivary metals data

We obtained salivary metals concentrations in children from Gatzke-Kopp et al 2023 (14) and report their methods here for clarity. Briefly, Gatzke-Kopp and colleagues used Inductively Coupled Plasma Optical Emission Spectrometry (ICP-OES) to measure the concentration of metals in aliquots of saliva that we matched to our own. Due to uneven sampling of salivary metals, we obtained data for lithium (N = 204), chromium (N = 237), copper (N = 237), manganese (N = 237), and zinc (N = 237).

### Metabolome sample preparation and mass spectrometry

We sent frozen saliva samples on dry ice to the West Coast Metabolomics Center (WCMC) at the University of California, Davis for sample preparation and data collection. Briefly, WCMC extracted 100 uL of saliva with 3:3:2 acetonitrile/isopropanol/water then evaporated the samples. WCMC ran 5 uL of sample through hydrophilic interaction chromatography (HILIC) then used an Agilent 6530 Quadrupole Time of Flight MS/MS mass spectrometer in both positive and negative ion modes to capture ion intensities. WCMC used published protocols to collect data, and ran all samples, quality control pool samples, and method blanks (Cajka and Fiehn 2016; Kind et al. 2018, 2009) in the same manner. We note that within each sample WCMC added stable isotope “internal control standards” (iSTDs) to the samples before injection to calibrate retention times. We received the data as an Excel file with metabolite names (for identified metabolites only), adducts, ion mode, retention times, mass/charge ratios (m/z), and peak heights for each metabolite (Dryad dataset (40)). We then assigned “chemical taxonomy” with the chemical classification software “ClassyFire” (Djoumbou Feunang et al. 2016).

### Metabolome data analyses

We used R (R Core Team 2018) for data analyses and manipulation. Before any analyses, we removed iSTD compounds from the dataset, as the WCMC added these compounds for quality control. We also filtered out ions with an average relative abundance of less than two-fold higher in samples or quality control samples as compared to blanks. We normalized the metabolites to their within-sample relative abundances and used these relative abundances for downstream analyses. We then used these data to tabulate Shannon diversity and Bray-Curtis indexes and visualized the data with nonmetric multidimensional scaling ordinations (NMDS) with “ggplot2” (Wickham 2009) and “patchwork” (41) with categorical variables self-reported on a questionnaire: “caregiver/child”, “smoking/nonsmoking” for caregivers, and “<1 ng/mL or >1 ng/mL” cotinine concentration for children as an estimate for ETS exposure (16). We tested these variables for statistical significance with Adonis (PERMANOVA with 999 permutations) in the R package “vegan” (42) for multivariate statistics and linear mixed effects models on individual metabolites with “lmerTest” (43) in R. We obtained salivary metals data from Gatzke-Kopp et al 2023 (14) on sample aliquots from the same original source, and refer to their publication for all relevant materials and methods to metals concentration analyses. Similarly, we assigned metals concentrations to tertile based on concentration within the entire subsample and ran Adonis tests on those data as above.

We used the R package “Hmisc” (Harrell 2019) to generate Spearman correlations between the relative abundances of metabolites and the concentrations of the biomeasures adiponectin, cotinine, CRP, and uric acid, along with each metal for children and cotinine for caregivers only, then corrected the resulting P-values for multiple comparisons via the Benjamini-Hochberg method. We also used distance-based Redundancy Analysis (db-RDA) in “vegan” to assess the contributions of continuous variables to microbiome variability and plotted the resulting ordinations with “ggplot2.”

Lastly, we performed stomatotype analysis (as in (18,44,45)) on the metabolomes of paired caregiver/child dyads using Jensen-Shannon distances and partitioning around medoid (PAM) clustering with the R packages “ade4” (46) and “cluster” (47). We estimated optimal cluster number by Silhouette values, then used lmerTest for univariate analyses between PAM clusters, adjusted p-values for multiple comparisons with the Benjamini-Hochberg method, and plotted the resulting PCoA ordination and boxplots of the data with “ggplot2” and “patchwork.” *Data availability:*

Tabulated mass-spectrometry metabolomics data are available on Dryad (40), and representative code used to analyze the data will be available on GitHub at github.com/jasonarothman/ECHO_90mo_metabolome.

## Results

We obtained metabolite profiles from both caregivers (N = 706) and children (N = 719), along with extraction method blanks (N = 153), and quality control pool samples (N = 157) (Dryad dataset (40)). From these metabolomes, we detected 2,881 unique metabolites, of which 281 could be identified, and filtering left us 2,046 total metabolites (both positive and negative ions) of which 210 were identified. Overall, the 104 most proportionally abundant metabolites comprised 50.02% of the relative abundance in our samples (Fig. S1), of which 1,409 were present at > 0.01% relative abundance. Likewise, identified metabolites only comprised 15.56% of the relative abundance of metabolites, while unknown compounds accounted for 84.45%. We used the ClassyFire chemical classifier to assign “chemical taxonomy” to each identified compound and plotted the relative abundances of the ten most proportionally abundant chemical superclasses as a stacked bar plot (Figs. 1 and S2).

**Figure 1:**
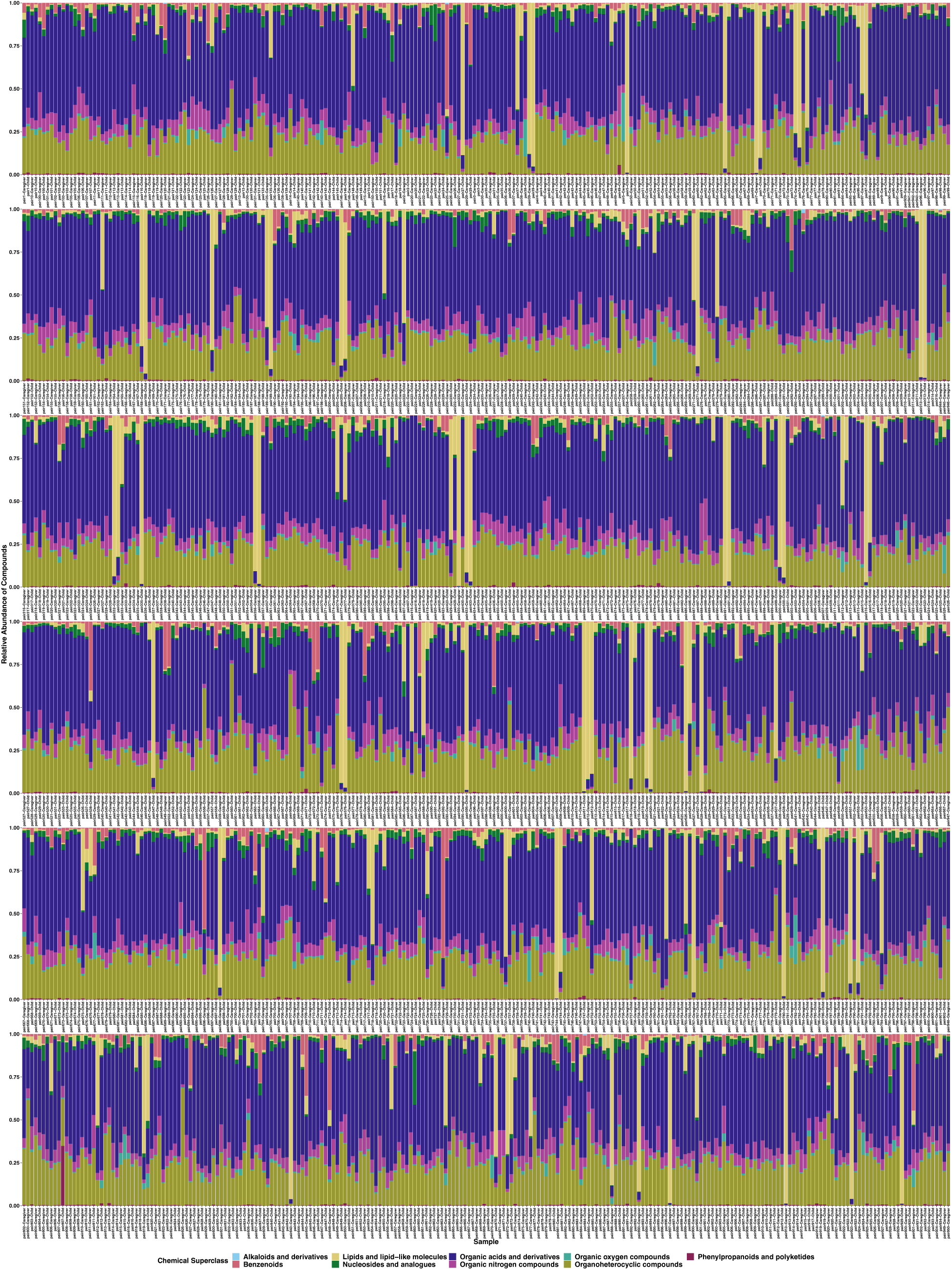
Stacked bar plot showing the relative abundances of the chemical superclass of 281 identified compounds across samples as assigned by ClassyFire. Color denotes the chemical superclass.

We paired 1426 samples into 672 caregiver/child dyads. We compared the diversity of all 2,046 metabolites (as measured by Bray-Curtis dissimilarities) and found that children and caregivers significantly differed (F = 37.2, R^2^ = 0.02, P < 0.001), and dyad explained most of the variation between our subjects (F = 1.8, R^2^ = 0.62, P < 0.001, Fig. 2a), while alpha diversity was not different between caregivers and children (H(_1_) = 1.6, P = 0.20). Then, we used linear mixed effects models with dyad as a random variable on metabolites at greater than 0.01% relative abundance and showed that 95 identified metabolites differed between caregivers and children (P_adj_ < 0.05, supplemental file SF1). Of these, 20 metabolites were highly significantly different between caregivers and children (P_adj_ < 2 x 10^-15^): Theophylline-, caffeine+, 4-imidazoleacrylic acid+/-, ribose-5-phosphate-, inosine-, agmatine+, 2-amino-1-phenylethanol+, and amino acids and their derivatives (Fig. 2, supplemental file SF1).

**Figure 2:**
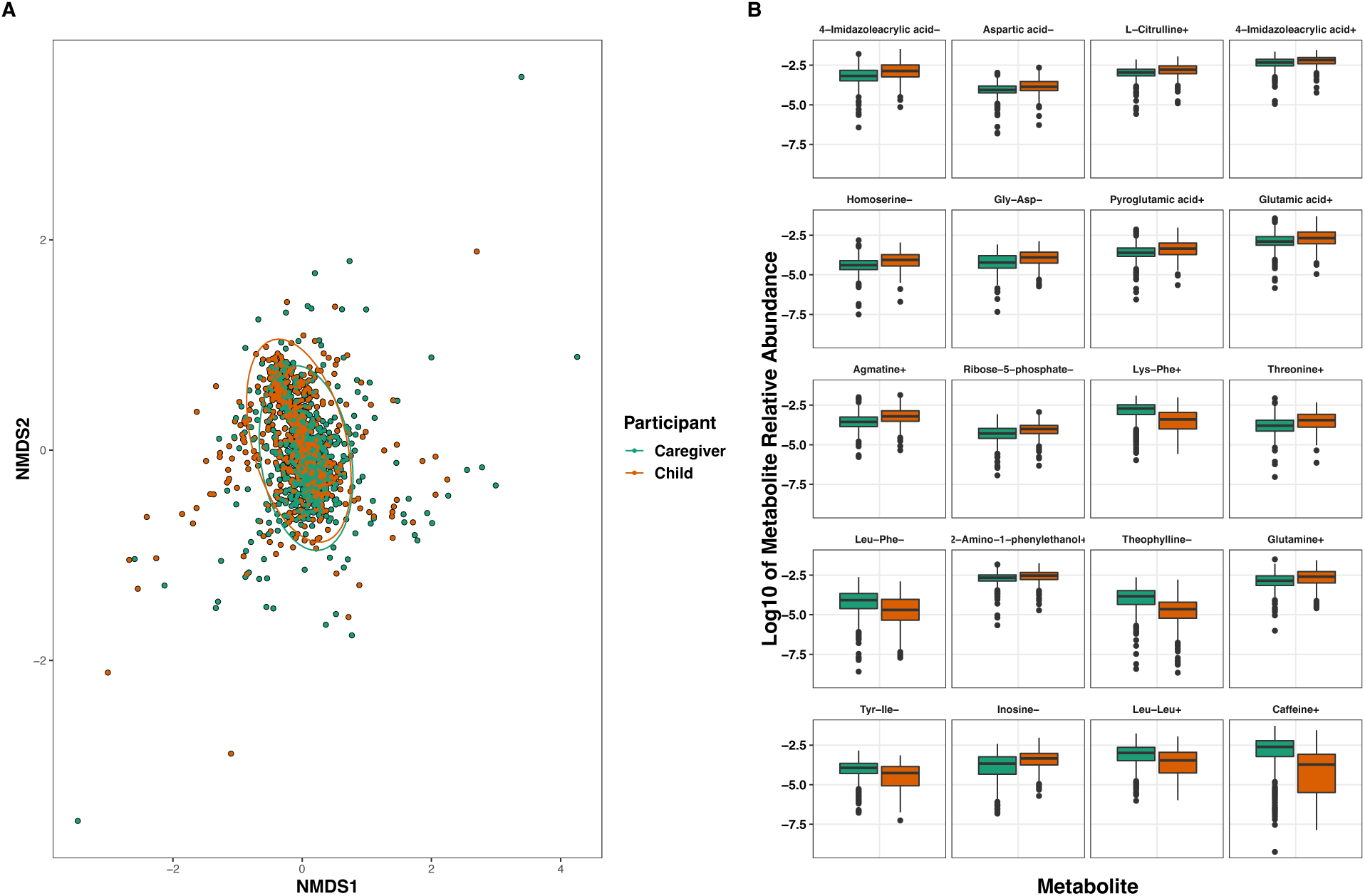
Comparisons of caregivers and children by A) NMDS ordination and B) linear mixed effects models of the relative abundances of the 20 most significantly differentially abundant metabolites (P_adj_ < 2 x 10^-15^). Dyad explained most of the variation between our subjects (F = 1.8, R^2^ = 0.62, P < 0.001 while categorical age explained approximately 2% of metabolome variation (F = 37.2, R^2^ = 0.02, P < 0.001). (+) or (-) denotes positive or negative metabolite ions.

We analyzed the metabolomes of paired caregiver/child dyads (N = 1344) through PAM clustering and observed that these metabolomes clustered into two overlapping groups of individuals based on Silhouette values (Silhouette = 0.12) and plotted the resulting PCoA ordination (Fig. 3a). Metabolite beta diversity differed between the two clusters (F = 189.6, R^2^ = 0.12, P < 0.001). We then ran linear mixed effects models on metabolites at greater than 0.1% relative abundance with dyad as a random variable to univariately compare the clusters and showed that 25 identified metabolites significantly differed between the clusters (P_adj_ < 0.05, Fig. 3b, supplemental file SF1). Of these, 20 metabolites had P_adj_ values of < 1 x 10^-6^: Hypoxanthine+, oxypuranol-, 2-amino-1-phenylethanol+, 2-(4-amino-1-piperidinyl)ethanol+, 4- imidazoleacrylic acid+, histamine+, nudifloramide+, phenylacetaldehyde B+, and amino acids and derivatives (including nonproteinogenic) (Fig. 3, supplemental file SF1).

**Figure 3:**
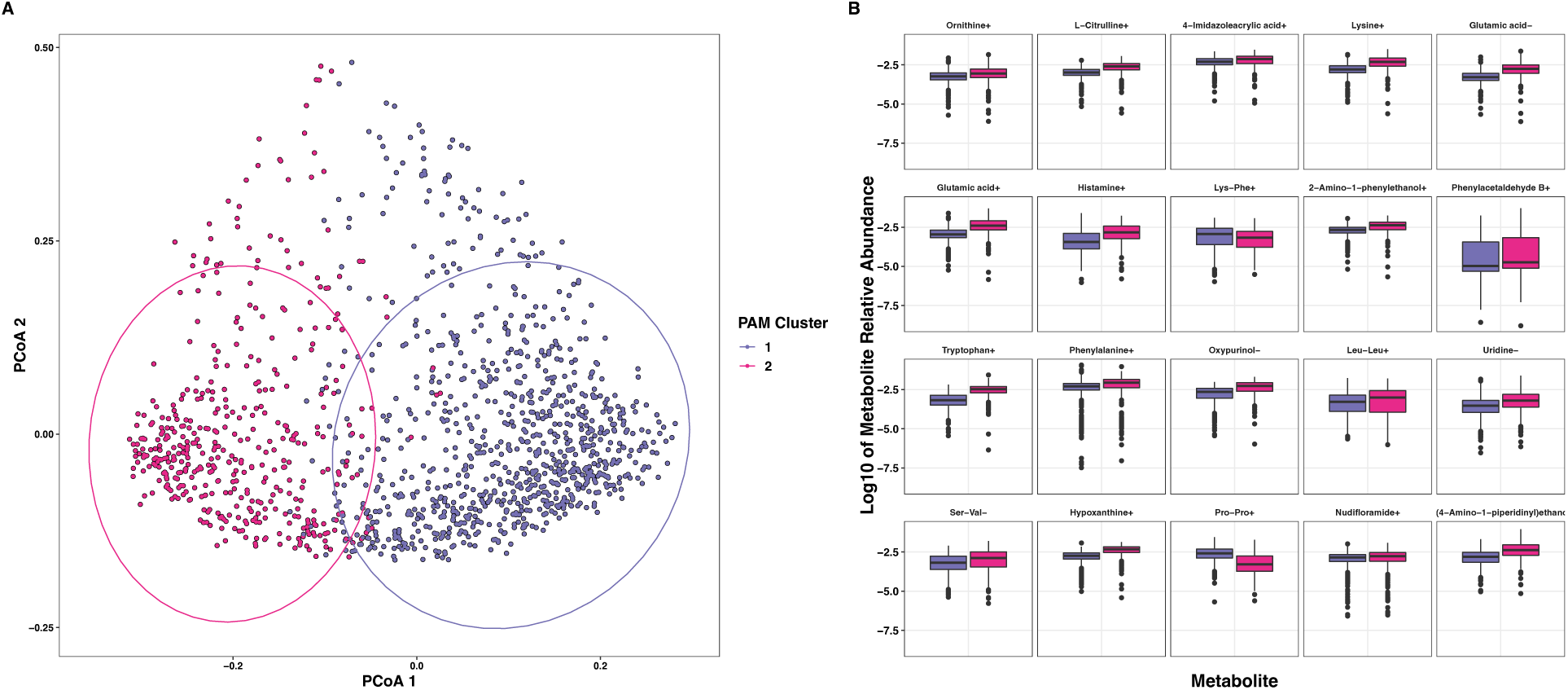
Analyses of the metabolomes of paired subjects by A) PAM clustering and B) linear mixed effects models of the relative abundances of the 25 most significantly differentially abundant metabolites (P_adj_ < 0.001). Metabolomes clustered into two overlapping groups, and only metabolites at greater than 0.1% relative abundance were analyzed by LMER. (+) or (-) denotes positive or negative metabolite ions.

We assessed the associations between concentrations of adiponectin, CRP, cotinine, and uric acid with the overall metabolomes of children (N = 538) through distance-based redundancy analysis (db-RDA). We observed that the four biomeasures cumulatively explained 5.3% of metabolome variation (Overall F = 7.5, P < 0.001), with each compound significantly associated with the metabolomes (P < 0.026 for each, Fig. 4, supplemental file SF1).

**Figure 4:**
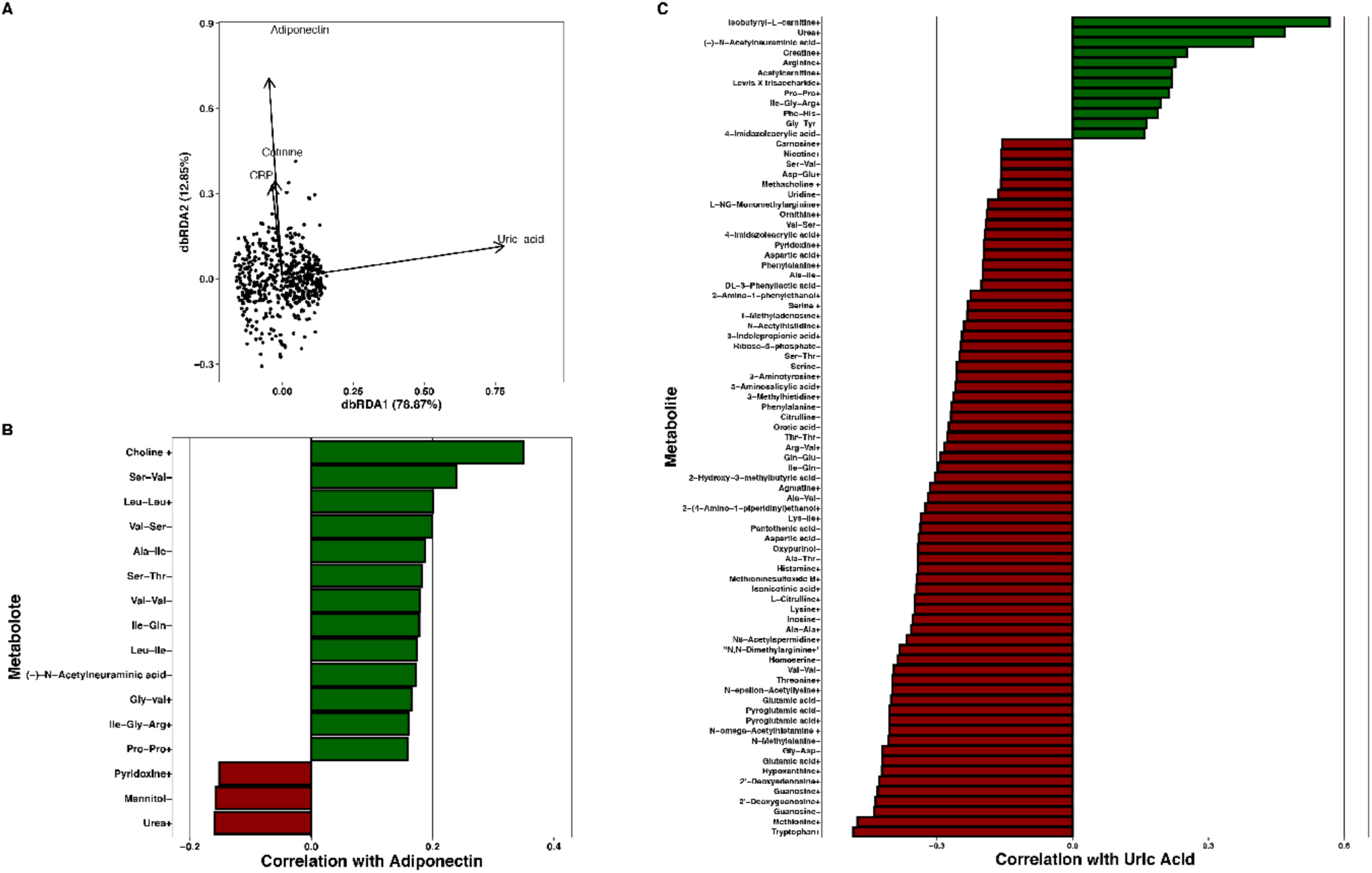
A) db-RDA analysis of children’s metabolomes with kit-assayed biomeasure concentrations overlayed as vectors. Overall, the four measured biomarkers significantly associated with metabolome separation (F = 7.5, P < 0.001), with each compound significantly associating with the metabolomes (P < 0.026 for each). B-C) Significant Spearman correlations (-0.15 < ρ > 0.15 shown only) of identified metabolites with kit-assayed adiponectin and uric acid. (+) or (-) denotes positive or negative metabolite ions.

We ran univariate Spearman’s correlations on metabolites at greater than 0.01% average relative abundance to find associations between specific compounds and each of the salivary biomeasures examined. We report the sample sizes for each analysis as each biomeasure had a different number of measurements. Concentrations of adiponectin were significantly correlated with 41 metabolites (N = 701, -0.16 < ρ < 0.35, P_adj_ < 0.05, Fig. 4, supplemental file SF1). CRP levels were significantly correlated with 22 metabolites (N = 616, -0.13 < ρ < 0.15, P_adj_ < 0.05, Fig. 4, supplemental file SF1). Lastly, uric acid concentrations were significantly correlated with 94 metabolites (N = 630, -0.49 < ρ < 0.57, P_adj_ < 0.05, Fig. 4, supplemental file SF1). We also ran these correlation tests on kit-measured cotinine concentrations and report those results with the smoking/nonsmoking analyses.

We compared the associations between ETS exposure and children’s (N = 714) and caregivers’ (N = 672) metabolomes separately because of the different routes of smoking exposure. We analyzed children’s metabolomes categorically classifying children by their kit-measured cotinine concentration (children’s cotinine levels <1 ng/uL and >1 ng/uL) but did not see a significant relations between ETS exposure and metabolic alpha (H(_1_) = 0.25, P = 0.61) or beta diversity (F = 1.1, R^2^ = 0.002, P = 0.29, Fig. 6a). We then ran generalized linear models on each metabolite and found that only nicotine+ was significantly different between groups (P_adj_ < 0.001, supplemental file SF1).

**Figure 6:**
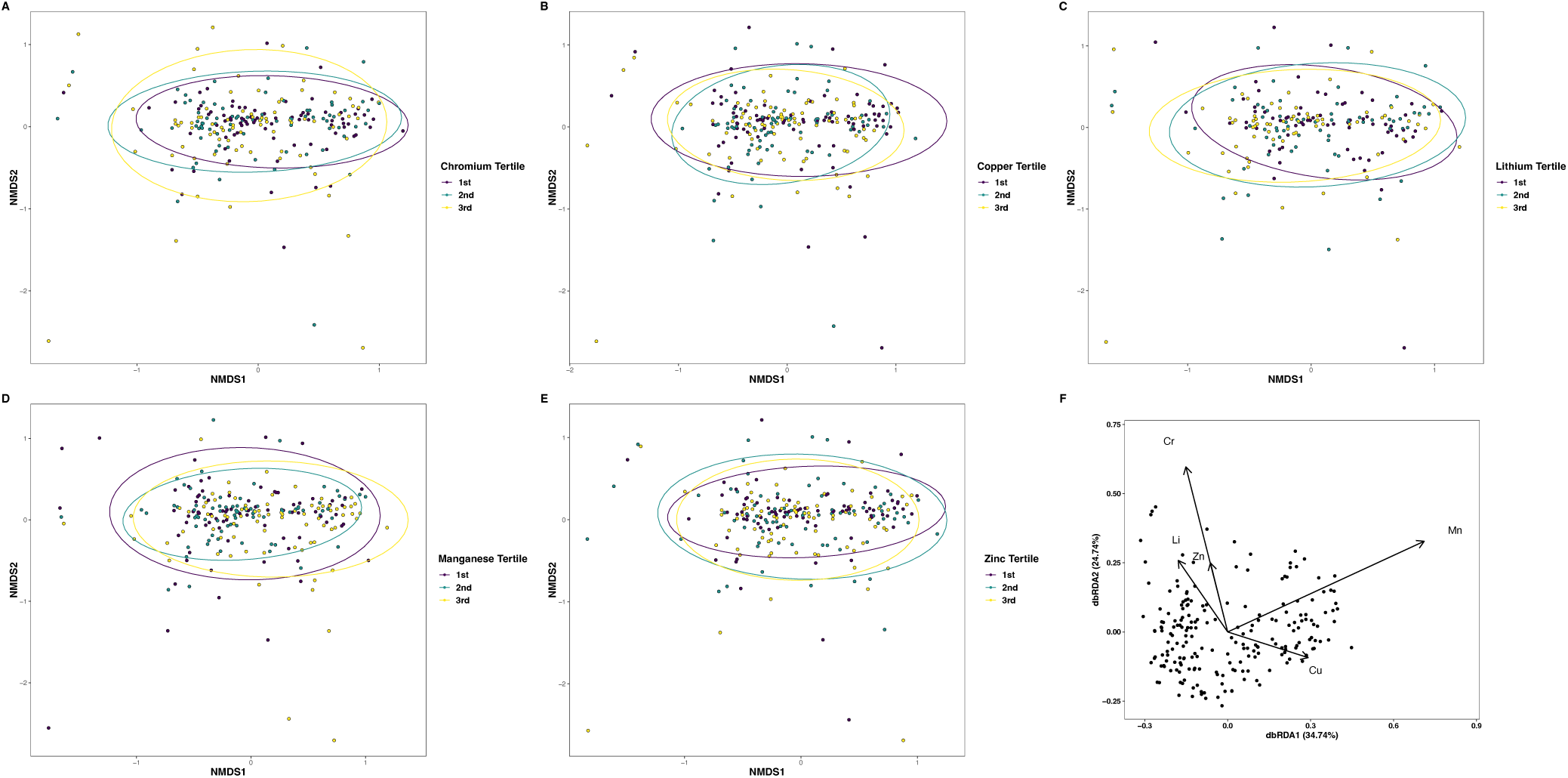
NMDS ordinations of salivary metal tertiles for A) chromium, B) copper, C) lithium, D) manganese, and E) zinc for each metal tested colored by tertile. Metals significantly associated with metabolome beta diversity ([Li: R^2^ = 0.01, F = 1.5, P = 0.044], Cr: [R^2^ = 0.01, F = 1.5, P = 0.039], Cu: [R^2^ = 0.02, F = 2.1, P = 0.004], Mn: [R^2^ = 0.02, F = 2.7, P < 0.001], Zn: [R^2^ = 0.01, F = 1.6, P = 0.038]). Panel F is a db-RDA of all metal concentrations. Metals explained 4.6% of metabolome variation (Overall F = 1.9, P < 0.001), and chromium, manganese, and copper contributed significantly to the variation, while lithium and zinc did not (Cr, Mn, Cu: P < 0.05, Li and Zn: P > 0.05).

As the intensity of ETS exposure likely causes different effects to the metabolome we ran Spearman correlations between kit-assayed cotinine and individual metabolites in children (N = 714). Cotinine was significantly correlated with 14 metabolites (-0.11 < ρ < 0.26, P_adj_ < 0.05, Fig. 5, supplemental file SF1).

**Figure 5:**
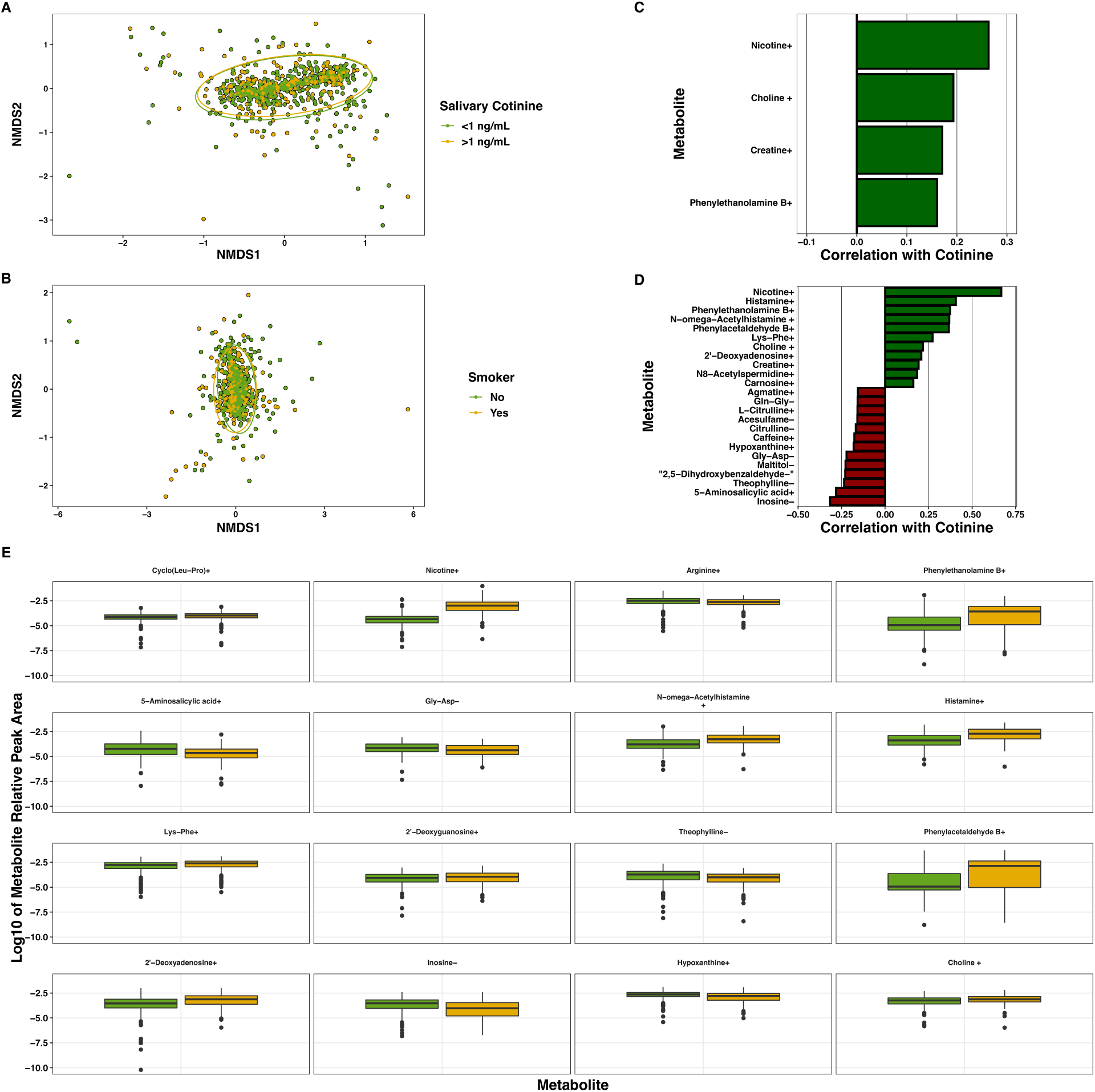
A) NMDS ordination of the metabolomes of children with salivary cotinine <1 ng/uL versus >1 ng/uL and B) caregivers who reported smoking or not. ETS exposure did not significantly alter the metabolome of children (F = 1.1, R^2^ = 0.002, P = 0.29), but did affect caregivers’ (F = 8.6, R^2^ = 0.01, P < 0.001. C & D) Significant Spearman correlations (P_adj_ < 0.05, (-0.15 < ρ > 0.15 shown only) of salivary cotinine and kit-measured cotinine in C) children or D) caregivers. E) Boxplots of the relative abundances of significantly different metabolites between smoking and nonsmoking caregivers as tested by generalized linear models (P_adj_ < 0.05, only P_adj_ < 0.01 shown). (+) or (-) denotes positive or negative metabolite ions.

We found that self-reported smoking status slightly related to the metabolome beta diversity of caregivers (F = 8.6, R^2^ = 0.01, P < 0.001, Fig. 5), but not alpha diversity (H(_1_) = 0.44, P = 0.51). We used generalized linear models on individual metabolites at greater than 0.01% relative abundance to find compounds that associated with caregivers’ smoking status. Only 22 identified metabolites significantly differed between groups: Nicotine+, 4-imidazoleacrylic acid+, phenylethanolamine B+, 5-aminosalicylic acid+, N-omega-acetylhistamine+, histamine+, 2’-deoxyguanosine+, theophylline-, phenylacetaldehyde B+, 2’-deoxyadenosine+, inosine-, hypoxanthine+, choline+, nudifloramide+, and amino acids and derivatives (Fig. 5, supplemental file SF1).

We ran Spearman correlations between individual metabolites and kit-measured cotinine in caregivers in the same fashion as above in children. Cotinine was significantly correlated with 39 identified metabolites (-0.32 < ρ < 0.67, P_adj_ < 0.05, Fig. 5, supplemental file SF1).

We measured the associations between the metals lithium (N = 204; average = 13.9 μg/L, range = 0.11 – 845.5 μg/L), chromium (N = 237; average = 7.3 μg/L, range = 0.07 – 22.5 μg/L), copper (N = 237; average = 29.0 μg/L, range = 0.23 – 844.2 μg/L), manganese (N = 237; average = 8.6 μg/L, range = 0.04 – 122.5 μg/L), and zinc (N = 237; average = 60.9 μg/L, range = 0.09 – 566.0 μg/L) measured in saliva and the metabolomes in children. As the metals were reported as concentrations (ug/L), we split the metals results into tertiles (i.e. 1^st^, 2^nd^, 3^rd^) to analyze the associations categorically. Salivary metal tertile for each element significantly associated with metabolome beta diversity ([Li: R^2^ = 0.01, F = 1.5, P = 0.044], Cr: [R^2^ = 0.01, F = 1.5, P = 0.039], Cu: [R^2^ = 0.02, F = 2.1, P = 0.004], Mn: [R^2^ = 0.02, F = 2.7, P < 0.001], Zn: [R^2^ = 0.01, F = 1.6, P = 0.038], Fig. 6), but not metabolome alpha diversity (P > 0.05 for each, Fig. 6). We then used db-RDA on the samples with measured concentrations of all metals (N = 204) and found that these metals explained 4.6% of metabolome variation (Overall F = 1.9, P < 0.001), but only chromium, manganese, and copper contributed meaningfully to this variation, while lithium and zinc did not (Cr, Mn, Cu: P < 0.05, Li and Zn: P > 0.05, Fig. 6, supplemental file SF1).

We also measured univariate Spearman correlations between metals concentration and individual metabolites at greater than 0.01% relative abundance. Lithium was significantly correlated with six metabolites (ρ < -0.21, P_adj_ < 0.05, Fig. 7, supplemental file SF1), chromium was significantly correlated with nine metabolites (ρ < -0.18, P_adj_ < 0.05, Fig. 7, supplemental file SF1), copper was significantly correlated with 42 metabolites (-0.46 < ρ < 0.44, P_adj_ < 0.05, Fig. 7, supplemental file SF1), manganese was significantly correlated with 52 metabolites (-0.47 < ρ < 0.38, P_adj_ < 0.05, Fig. 7, supplemental file SF1), and zinc significantly correlated with 29 metabolites (-0.43 < ρ < 0.31, P_adj_ < 0.05, Fig. 7, supplemental file SF1).

**Figure 7:**
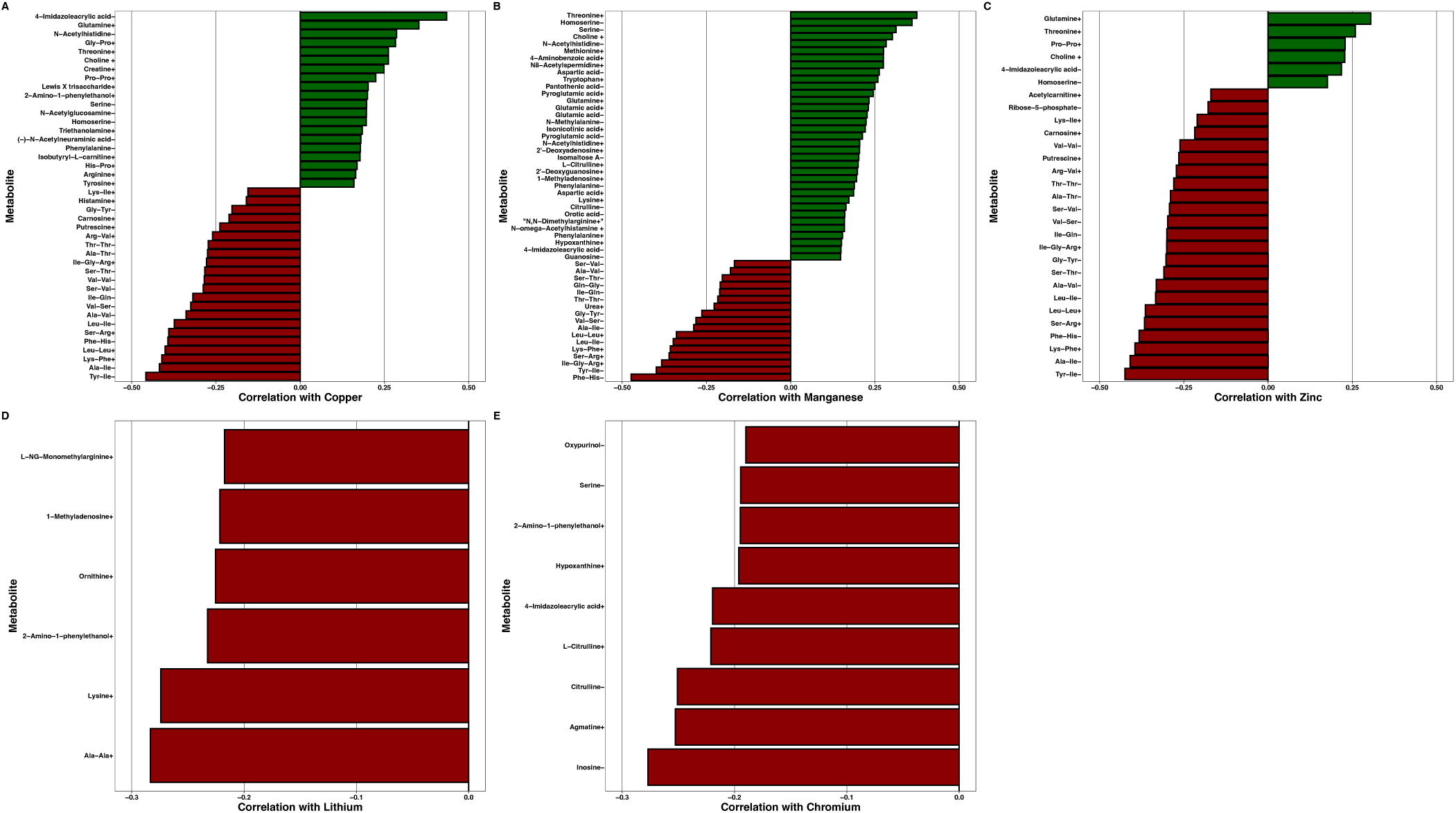
Significant Spearman correlations between metals concentration and individual metabolites at greater than 0.01% relative abundance (P_adj_ < 0.05) for A) copper, B) manganese, (C) zinc, D) lithium, and E) chromium. (+) or (-) denotes positive or negative metabolite ions.

## Discussion

Through our exploratory analyses of a large-scale sample from the 90-month assessment of the Family Life Project cohort (36), we were able to assay the salivary metabolome and its associations with biomeasures relating to antioxidant potential, environmental tobacco smoke (ETS) exposure, and systemic inflammation. Most of the variance in salivary metabolites were explained by family dyad, supporting the view that home environments are incredibly impactful on people, and the closeness of families often result in similar metabolisms and health measures. We also find that a small but significant proportion of the variance is explained by how children’s and adults’ metabolomes differed overall (22–26). Extending this concept further, subjects generally clustered into two overlapping metabolomic groups, suggesting that people fall into sub-groups with shared features, although we are unable to discern exactly what is causes this grouping (1). We observed significant but minor shifts in smokers’ metabolomes, but not in their children, indicating that primary ETS exposure can affect salivary metabolites and may be associated with increased inflammation and polyamine turnover (4,31,48,49). We also show that kit-based biomeasures correspond well with compounds intermediate in their respective biochemical pathways, suggesting both kit-based and mass-spectrometry-based approaches are useful to assaying metabolism. Lastly, salivary metals associated with altered children’s metabolomes, and were often anticorrelated with putative proteolysis products, suggesting that environmental metals may affect metabolism (50), and that saliva is a useful biofluid in which to assay metals (14). Collectively, our study represents a large-scale investigation into the metabolism of families and how these compounds may relate to the cohabitating individuals.

Our study shows that caregiver/child dyad explained most of the salivary metabolome variation, supporting others’ similar findings in specific metabolites (22–25), but previously unknown in large-scale studies measuring the overall metabolome. Many factors are known to affect metabolism, such as age, sex, diet, and increasingly, familial environment, and our work adds an important dimension to this metabolic variability (1,25,27). By applying untargeted metabolomics, we can start to unravel the complex interactions that family and social structure has on the metabolism of individuals. Even though child/caregiver dyad explained the vast majority of metabolic variation, several metabolites’ relative abundance differed between children and caregivers - many of which are “lifestyle,” medicinal, or environmental compounds/metabolites. Compounds such as nicotine, caffeine, theophylline, phenylacetaldehyde, acetaminophen, and salicylic acid were all higher in adults, likely due to diet or medication (4). Similarly, adults generally had higher levels of dipeptides (suggesting proteolysis or oxidative damage), while children had more metabolites basal or intermediate to chemical pathways, perhaps due to children’s’ higher basal metabolic rate and growth. As amino acid metabolism has been shown to change with age (4) and differs across populations (1), more research is needed to understand amino acid metabolism associations with medical conditions or normal development. We also note that large proportions of our data consist of unidentified metabolites and are likely important to discerning child from caregiver, but we are unable to classify these compounds. We appreciate that the magnitude of these differences, while significant, are slight, further stressing the impact and importance of family (relatedness and/or cohabitation) on metabolism.

Unsupervised clustering of large-scale, multidimensional metabolomics data has been used to group diseased and healthy individuals by common metabolic patterns and provide insights into their underlying physiology (1,9,30,51–53). In our study, subjects’ metabolomes tended to cluster into two overlapping groups which did not obviously correspond to any of our metadata categories (i.e. sex, age, smoking, state of residence, etc.). These clusters were somewhat separated by free dipeptides, a Lewis X trisaccharide (Le^X^) and intermediates of the urea cycle (i.e. citrulline, arginine, and ornithine), but mainly differed by the proportion of unidentified, higher-abundance metabolites, and we note that the overall differences between clusters were relatively minor as seen before (1). Previous cluster-based metabolomics studies have shown that differences in amino acid metabolism can cause cluster separation (1,9,30), and oral microorganisms impact the concentrations of these metabolites (1,15), so future studies should incorporate “multi-‘omic” approaches to understanding the chemical and microbial ecology of the mouth. Likewise, urea cycle metabolites were important in cluster separation. Intermediates of the urea cycle have been implicated as potential indicators for diverse maladies, such as hypertension (54), breast cancer (30), severe inflammation (55) and urea cycle disorders (56), suggesting that untargeted metabolomics and cluster analyses are useful to find compounds that may serve as biomarkers for disease. While the clusters largely separate in ordination space, we note that there is substantial overlap, indicating variability and noise in our data, which may be affecting our interpretations. Similarly, as we are unable to pinpoint exactly what is driving metabolome clustering, our results may be influenced by diet or other personal choices that are not reflected in our metadata (1,2,4,15,57,58).

Adiponectin, CRP, and uric acid significantly associated with altered metabolomes in children, with uric acid explaining the most variance. As the endpoint in purine degradation (59), uric acid was (as expected) strongly anticorrelated with intermediates of purine metabolism. Conversely, acylcarnitines were correlated with uric acid, which may indicate consumption of a high-fat/high-protein diet and subsequent lipid accumulation (60), and an increase in creatine with uric acid may also be due to protein intake and subsequent purine metabolism or antioxidant activity (61,62). We found many positive correlations between free dipeptides and adiponectin – a molecule predicted to modulate inflammation and oxidative stress (13,63). As dipeptides are thought to be biomarkers for proteolysis (64) and may have antioxidant activity (65) the correlations of adiponectin and dipeptides could indicate infections or inflammation in children’s mouths, and so may serve as a useful salivary biomeasure for immune activity. Lastly, CRP had very slight associations with oral metabolomes and individual metabolites, suggesting that while CRP is a useful biomarker for systemic inflammation, it is also nonspecific (66), and may not associate well with metabolic pathways in saliva (15). Collectively, these results suggest that the kit-based biomeasures capture useful physiological markers for their respective pathways or functions, and indicate that saliva is a worthwhile biospecimen for assaying these measures (10).

Environmental tobacco smoke (ETS) exposure is a major, worldwide cause of morbidity and has been shown to alter human metabolism (4,31,48). Smoking minorly affected caregivers’ overall metabolomes but exposure to ETS did not alter children’s – likely because children were only exposed passively and likely did not receive the same nicotine dose as active smokers. In both children and caregivers, several metabolites were correlated with kit-assayed cotinine (the major final metabolite of nicotine), including phenylethanolamine, phenylacetaldehyde, choline, creatine, N8-acetylspermidine, and unsurprisingly nicotine, along with a strong apparent histamine response in caregivers. As an agonist of acetylcholine receptors, nicotine has been shown to increase acetylcholine demand (and therefore choline catabolized from phospholipids) (67), which may be driving our perceived proportional increase in salivary choline (68), along with increasing phenylethanolamine N-methyltransferase activity (possibly observed in our study as increased phenylethanolamine) (69). Likewise, smoking reduces creatine kinase activity (converts creatine to phosphocreatine), which may be causing higher levels of creatine with increasing nicotine concentrations (70,71). In regard to N8-acetylspermine – a compound indicative of polyamine turnover and associated with vascular pathologies (49,72) – again we observed a correlation with nicotine. Lastly, histamine was markedly correlated to cotinine concentration, suggesting that histamine production, mast cell activations, and inflammation are affected by nicotine use in caregivers (73,74). While the overall metabolomic associations with self-reported smoking status and cotinine concentration were minor, the above individual correlations indicate that saliva metabolomics are useful to find potential biomarkers that may indicate altered physiology related to tobacco smoke exposure (4,31,48).

Salivary metal concentrations were slightly associated with altered child metabolomes, suggesting that metals consumption/exposure potentially affects metabolism (32,75–78). As the metals cooccurred in the samples, we analyzed them simultaneously, and show that there are likely synergistic associations with the metabolome, which shifts the individuals’ overall metabolism (77). When specifically considering zinc, copper, and manganese, metals concentration were often anticorrelated with free dipeptides – potential markers for proteolysis and oxidative stress – suggesting that these metals may act as antioxidants reducing the body’s need to produce its own dipeptide antioxidants (64,65). Conversely, there were positive associations between these metals and free amino acids, which may indicate a shift toward complete protein degradation or proteosome activation (50,78). Taken together, these results suggest that metals are involved in protein metabolism or that metals may be acting as or along with antioxidants, but more research into oral metabolomics is needed. Still, our work suggests that metals exposure and the metabolome interact, and more studies should incorporate these types of analyses to better understand human metabolism.

### Conclusions

Metabolomics allows for the simultaneous detection of thousands of chemicals and can provide insights into the complex biochemical activity of human metabolism. Our study suggests that there are both large-scale patterns and subtle differences in the salivary metabolome between populations and within families, and that cohabitation likely affects metabolism. Taken together, metabolomics and kit-based biomeasure analyses indicate that tobacco smoke affects primary users’ metabolism, and that there are several putative biomeasures for antioxidant potential, tobacco smoke ETS exposure, and systemic inflammation along with metals concentrations, that can be studied for further use in understanding the biochemistry of environmental exposures and stress. Furthering our inquiry, we suggest that future research investigate the interactions between the oral microbiome and metabolome, and that “multi-omic” approaches be applied to family-based or large-populational studies to understand the complex microbes and molecules that underly human health.

## Acknowledgements

This study is part of the Family Life Project (FLP). We thank Kaitlin Smith and Tatum Stauffer for technical assistance with salivary biospecimen testing and the Family Life Project (FLP) Investigators. We would like to express our gratitude to the families, children, and teachers who participated in this research and to the FLP research assistants for their hard work and dedication. This research was supported by The National Institute of Child Health and Development (NICHD) Award #P01HD039667, and #R21HD061649, the Environmental influences on Child Health Outcomes (ECHO) program, Office of The Director, National Institutes of Health (NIH) Awards #1UG3OD023332, #1UH3OD023332, and Hewitt Foundation for Biomedical Research and National Institute of Aging T32 (#AG00096-40) postdoctoral fellowships to JAR.

