## Supplemental Information for "Salivary metabolomics in the family environment: A large-scale study investigating oral metabolomes in children and their parental caregivers"

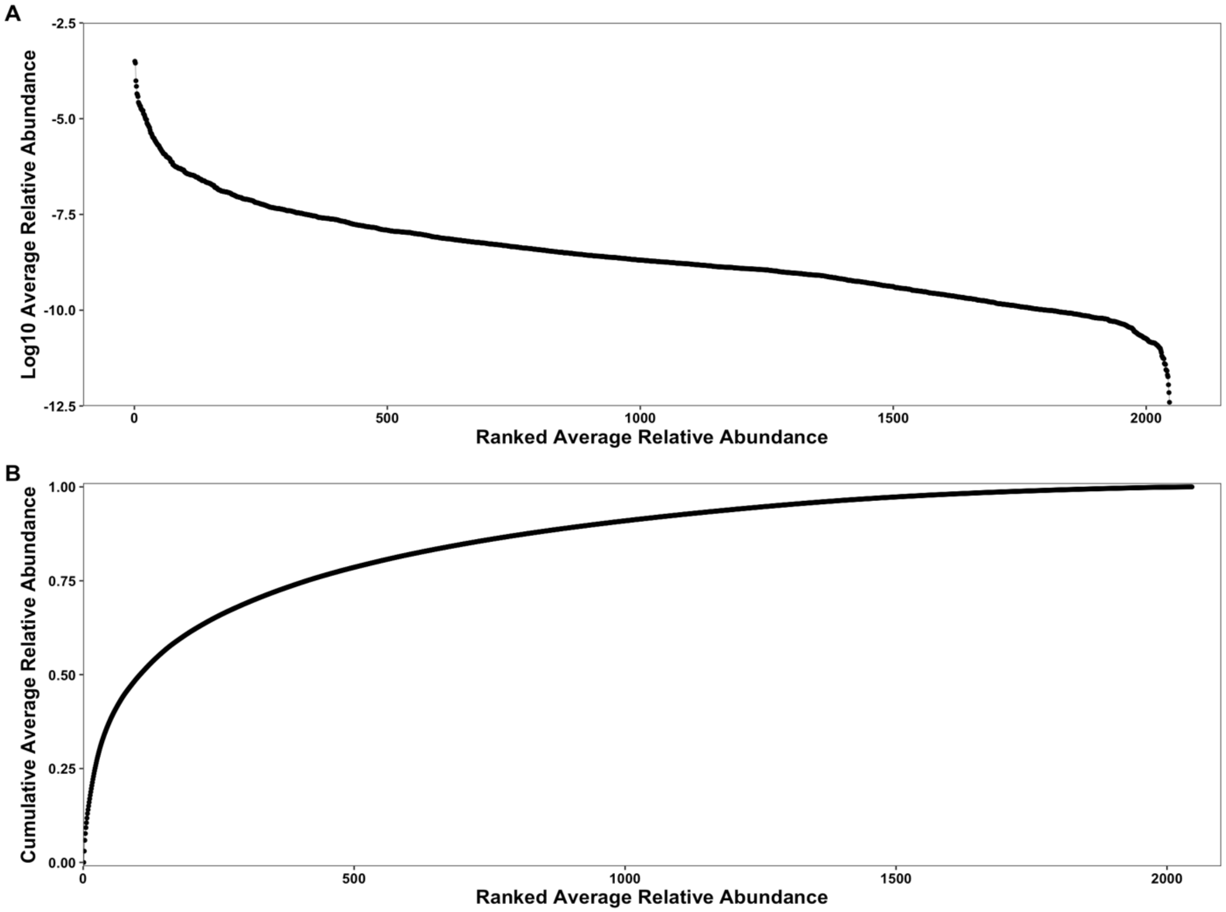


Figure S1: A) Rank abundance curve of the $log_{10}$ transformed average relative abundances of metabolites across all samples. B) Accumulation curve of the ranked average relative abundance of metabolites across all samples.

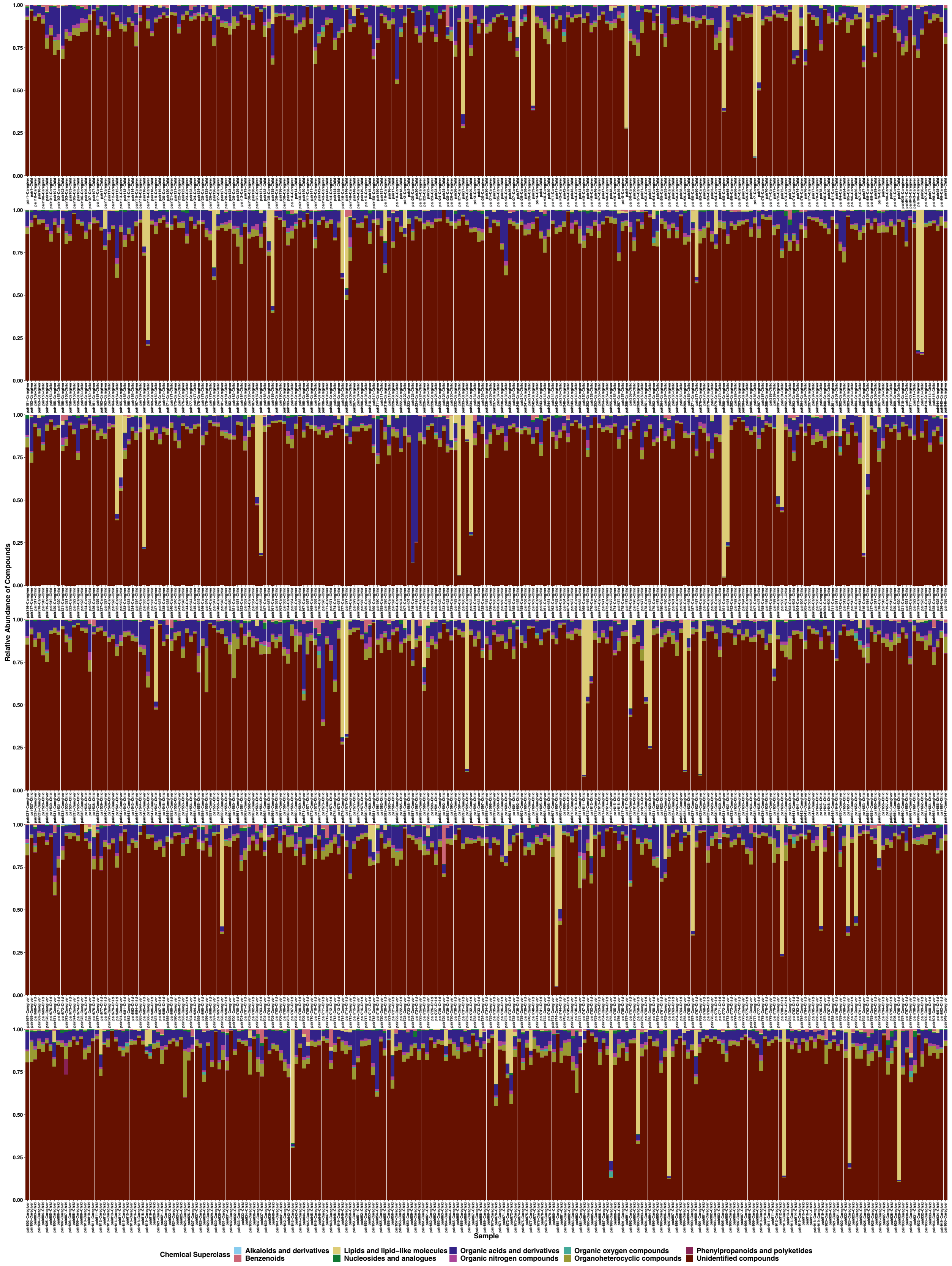
Relative Abundance of Compounds
1.00
0.75
0.50
0.25
0.00
Sample
Chemical Superclass
Alkaloids and derivatives
Benzenoids
Lipids and lipid−like molecules
Nucleosides and analogues
Organic acids and derivatives
Organic nitrogen compounds
Organic oxygen compounds
Organoheterocyclic compounds
Phenylpropanoids and polyketides
Unidentified compounds

Figure S2: Stacked bar plot showing the relative abundances of the chemical superclass of all compounds across samples either as assigned by ClassyFire or unidentified. Color denotes the chemical superclass.

Supplemental file SF1: Relevant statistical information for all Spearman correlation tests and linear models. Each worksheet tab contains information for individual tests. “Padj” denotes p-values adjusted for multiple comparisons.
